# A rapid ionic liquid-based DNA extraction method for molecular diagnostics of urinary tract infections

**DOI:** 10.1101/2025.09.30.679528

**Authors:** Johanna Kreuter, Lena Piglmann, Katarina Priselac, Roland Martzy, Michael Ante, Dominik Walter, Ildiko-Julia Pap, Barbara Ströbele, Andreas H. Farnleitner, Georg H. Reischer, Claudia Kolm

## Abstract

Rapid and reliable DNA extraction from urine is a critical bottleneck in advancing molecular diagnostics for urinary tract infections (UTIs) in both centralized and decentralized settings. Here, we present an ionic liquid-based DNA extraction method (IL-DEx) that enables recovery of bacterial DNA from urine samples in under 30 minutes using minimal equipment and no hazardous chemicals. IL-DEx was benchmarked against a widely used commercial kit (QIAamp DNA Mini Kit, QIAGEN) using reference strains, clinical isolates, and spiked urine samples. For gram-negative bacteria, IL-DEx achieved comparable DNA yields (47–102% relative efficiency), while recoveries from gram-positive bacteria were lower (0.7–8%) but sufficient for downstream detection. Quantitative PCR (qPCR) revealed linear DNA recovery across five to six orders of magnitude (10^8^–10^2^ CFU/ml, R^2^ >0.99), with detection limits of ∼10^2^–10^3^ CFU/ml for gram-negatives and ∼10^3^–10^4^ CFU/ml for gram-positives using 1 ml urine. Clinical evaluation with 13 patient urine samples (ten culture-positive, three culture-negative) demonstrated that IL-DEx reliably enabled pathogen detection by qPCR and full-length 16S rRNA gene sequencing (Oxford Nanopore). Performance was comparable to three other extraction methods tested head-to-head, including the QIAamp DNA Mini Kit (QIAGEN), the MagaZorb DNA Mini-Prep Kit (Promega), and a phenol-chloroform extraction method. These findings establish IL-DEx as the first ionic liquid-based approach evaluated for DNA recovery from clinical urine samples, providing a fast, simple, and low-cost method suitable for integration into molecular workflows for UTI diagnostics across diverse laboratory and clinical settings.

**Importance:** Urinary tract infections (UTIs) are among the most common infections worldwide and a major driver of antibiotic use. Rapid and accurate diagnosis is critical to guide therapy, reduce inappropriate antibiotic prescriptions, and improve patient outcomes. While molecular diagnostics can drastically reduce time to identify uropathogens, their implementation remains constrained by upstream DNA extraction – a step that is often laborious, cost-intensive, or incompatible with rapid diagnostic workflows. We developed a fast, simple, and low-cost DNA extraction method (IL-DEx) that uses an ionic liquid and magnetic beads to recover bacterial DNA directly from urine. IL-DEx eliminates hazardous reagents and complex equipment while delivering performance comparable to established extraction kits. By streamlining this critical pre-analytical step, IL-DEx enables faster molecular diagnostics and broadens access to modern UTI testing. Its simplicity and robustness position it as a valuable tool for improving diagnostic speed, antimicrobial stewardship, and patient care across healthcare settings.

## Introduction

Urinary tract infections (UTIs) are among the most prevalent infections worldwide, affecting more than 400 million individuals annually in both community and healthcare settings [1]. High-risk populations include infants, pregnant women, the elderly, catheterized patients, and individuals with chronic conditions such as diabetes. Nearly half of all women experience at least one UTI in their lifetime [2]. Rising global incidence [3], substantial healthcare costs [4, 5], and increasing antimicrobial resistance (AMR) among uropathogens highlight the need for timely and accurate diagnosis to guide therapy, reduce empirical antibiotic use and mitigate the spread of AMR [6, 7].

Clinically, UTIs present with a broad spectrum of etiologies and severities, ranging from uncomplicated lower tract infections, such as cystitis, to severe upper tract infections like pyelonephritis, which can progress to urosepsis or septic shock if not promptly treated [8, 9]. Uropathogenic *Escherichia coli* is the main causative agent, responsible for 65–75% of cases, followed by *Klebsiella pneumoniae, Proteus mirabilis, Pseudomonas aeruginosa, Enterococcus* spp. and *Staphylococcus* spp., depending on patient demographics and healthcare settings [10]. Current diagnostics rely primarily on urine culture, which requires 18-30 hours for species identification and an additional 18-24 hours for antimicrobial susceptibility testing [11]. While bacterial growth of 10^4^-10^5^ CFU/ml is usually considered indicative of a UTI, lower diagnostic thresholds (10^2^-10^4^ CFU/ml) can be clinically relevant in specific populations such as men, pregnant women or catheterized individuals [12, 13]. However, diagnosis is not always straightforward. Despite being the clinical gold standard, urine culture suffers from limited diagnostic sensitivity (e.g., ∼60% for detecting acute UTI [14]) and often fails to detect fastidious or polymicrobial infections [15-17].

These limitations have spurred the development of molecular diagnostic tools that provide rapid, sensitive, and often species-level detection of uropathogens and resistance genes directly from urine [18]. Emerging platforms include real-time multiplexed PCR assays [18, 19] and isothermal amplification assays such as loop-mediated isothermal amplification (LAMP), which are being explored for decentralized testing [20-22]. Likewise, both targeted (e.g., 16S–23S rRNA gene) [14, 23] and untargeted (shotgun metagenomics) [24] next-generation sequencing approaches are gaining traction as sequencing costs decline and bioinformatics tools mature. For instance, portable sequencers such as the MinION (Oxford Nanopore Technologies) have demonstrated near real-time identification of pathogens and resistance genes in urine in under five hours [25].

Despite these advances, all molecular platforms remain dependent on upstream nucleic acid extraction – a critical yet often overlooked step that impacts analytical performance, cost, and turnaround time [26]. Conventional extraction methods are often labor-intensive, expensive or poorly suited for rapid or decentralized workflows. To address this, we previously developed a simplified ionic liquid-based extraction protocol that combines chemical lysis with magnetic bead-based purification [27]. The method uses 1-ethyl-3-methylimidazolium acetate ([C_2_mim][OAc]) to achieve simultaneous bacterial cell lysis and nucleic acid binding, enabling DNA and RNA recovery from periopathogenic bacterial cultures within 30 minutes [27]. However, its performance on uropathogens and in urine – a matrix characterized by variable microbial loads, host DNA background, and inhibitory compounds – remained untested.

In this proof-of-concept study, we adapted and evaluated the ionic liquid-based extraction method (IL-DEx) for recovery of bacterial DNA from urine to support rapid molecular detection of uropathogens. IL-DEx was tested across a range of diagnostically relevant specimen types, including (i) cultured reference strains and clinical isolates, (ii) spiked artificial and native urine from healthy donors, and (iii) clinical urine specimens from patients with suspected UTIs. Benchmarking was performed against a widely used commercial kit (QIAamp DNA Mini Kit, Qiagen) and, for clinical specimens, extended to include a second commercial kit (MagaZorb DNA Mini-Prep Kit, Promega) and a traditional phenol-chloroform extraction method. Through this comparative framework, we aimed to assess the analytical performance and diagnostic utility of IL-DEx for integration into molecular workflows for UTI diagnostics.

## Methods

### Bacterial strains and isolates used in this study

This study included seven clinically relevant bacterial species commonly found in UTIs: *Escherichia coli* (type strain NCTC 9001; clinical isolate IHM 5531), *Pseudomonas aeruginosa* (reference strain NCTC 10662; isolate IHM 5716), *Klebsiella pneumoniae* (type strain DSM 30104; isolate IHM 5571), *Proteus mirabilis* (type strain DSM 4479; isolate IHM 5533), *Enterococcus faecalis* (type strain DSM 20478; isolate IHM 5490), *Enterococcus faecium* (type strain DSM 20477; isolate IHM 5458), and *Staphylococcus saprophyticus* (type strain DSM 20229; isolate IHM 5493). Isolates were obtained from urine samples of patients with confirmed or suspected urinary tract infections at University Hospital St. Pölten (Austria).

Gram-negative bacteria were cultured overnight in lysogeny broth (Merck, Germany) at 37 °C, with shaking at 170 rpm. Gram-positive bacteria were grown under identical conditions in tryptic soy broth (Merck). Overnight cultures were diluted in fresh medium to an optical density at 600nm (OD_600_) of 0.1 and incubated to reach an OD_600_ of ∼1.0 prior to use in downstream experiments.

### Experiments with pure bacterial cultures

Cells from actively growing cultures of *E. coli, P. aeruginosa, K. pneumoniae, P. mirabilis, E. faecalis, E. faecium*, and *S. saprophyticus* were harvested at an OD_600_ of ∼1.0. Cultures were centrifuged for 5 min at 4000 x g, washed once and resuspended in isotonic saline solution (0.9% NaCl), to the same OD_600_. For each strain, 10 µl of the resulting cell suspension were used for DNA extraction with the IL-DEx and the QIAamp DNA Mini Kit (QIAGEN, Germany). Extractions were performed in triplicate for each method. Negative controls using saline were included to monitor for contamination.

Total cell count per millilitre (TCC/ml) were determined by epifluorescence microscopy. For this, bacterial cell suspensions were fixed with sterile-filtered paraformaldehyde (final concentration 0.8%) overnight at 4 °C and then filtered through 0.2-µm Anodisc 25 filters (Whatman, Germany). Filters were stained with SYBR Gold (30 µl of 1:400 dilution in sterile deionized water; Invitrogen, Germany), incubated in the dark for 15 min, rinsed three times with sterile Milli-Q water, and air-dried in the dark. Filters were mounted on microscope slides using anti-fade mounting solution (Citifluor). Slides were examined with immersion oil using a Nikon Eclipse Ni microscope equipped with a Nikon DS-Qi2 camera at 400x or 1000x magnification (Ex ∼470, Em ∼515). TCC/ml were calculated based on the average number of cells counted at 10 randomly selected positions on the filter (10×100 mm^2^) and the volume of the sample filtered.

### Spiking experiments with artificial urine and urine from healthy donors

Actively growing cultures of *E. coli* NCTC 9001, *P. aeruginosa* NCTC 10662, *K. pneumoniae* DSM 30104, *P. mirabilis* DSM 4479, *E. faecalis* DSM 20478, *E. faecium* DSM 20477, and *S. saprophyticus* DSM 20229 with an OD_600_ of ∼1.0 were harvested by centrifugation (5 min, 4000 x g), washed, and resuspended in saline to an OD_600_ of ∼2.0. From each suspension, six tenfold serial dilutions (from 1:10 to 1:10^6^) were prepared. Each dilution and the undiluted cell suspension were spiked into artificial urine (Sigmatrix Urine Diluent, Sigma-Aldrich, USA) at a 1:10 ratio (v/v). A 1 ml aliquot of each spiked sample was centrifuged at 10000 x g for 10 min at 4 °C, and the pellet was subjected to DNA extraction using both IL-DEx and the QIAamp DNA Mini Kit (QIAGEN). Unspiked artificial urine was processed in parallel as a negative extraction control. All extractions were performed in triplicate for each dilution and method. For bacterial quantification, the three highest dilutions were used to determine colony forming units per ml (CFU/ml).

CFU/ml values were obtained by plating the respective cell dilution on lysogeny broth agar (Merck) in triplicate and incubating at 37 °C overnight. The number of colonies was counted and used for the calculation of CFU/ml. As negative control isotonic saline solution was plated.

Spiking experiments with urines from healthy donors were performed similarly. Freshly cultivated *E. coli* or *E. faecalis* cells were spiked to an OD_600_ of ∼0.02 into native urine samples from healthy donors. A 1 ml aliquot of each spiked and unspiked urine was centrifuged at 10000 x g for 10 min at 4 °C, and the resulting pellet was subjected to DNA extraction using both IL-DEx and the QIAamp DNA Mini Kit. To assess the DNA content of the spike, the cell suspension was also spiked into saline and processed in parallel. Negative extraction controls with only saline were included. All extractions were performed in triplicate per method.

### Clinical specimens

Thirteen clean-catch midstream urine samples from patients with suspected UTI were processed for comparative analysis of extraction methods: ten were urine culture-positive (>10^4^ CFU/ml) and three were culture-negative (< 10^4^ CFU/ml). Samples were collected in sterile containers and stored at 4 °C for no longer than five days prior to DNA extraction (see Table 1 and Supplementary I, Fig. S4). Ethical approval was not required, as all clinical samples used were archived coded remnant samples provided after completion of routine diagnostic testing for UTI. The study was conducted anonymously; no patient data was accessed or processed.

**Table 1.** Clinical metadata of urine samples for extraction and analysis.

| Sample No. | Urine appearance | Determined bacterial count by urine culture | UTI suspected | Identified uropathogen by MALDI-TOF | Storage time until extraction |
| --- | --- | --- | --- | --- | --- |
| 1 | Clear | $>10^4$ | Yes | *Citrobacter koseri* | 5 days |
| 2 | Bloody | $>10^4$ | Yes | *E. coli* | 5 days |
| 3 | Bloody, turbid | $>10^4$ | Yes | *E. coli* | 4 days |
| 4 | Clear | $>10^4$ | Yes | *E. coli* | 4 days |
| 5 | Slightly turbid | $>10^4$ | Yes | *E. coli* | 5 days |
| 6 | Turbid | $>10^4$ | Yes | *E. coli* | 4 days |
| 7 | Slightly turbid | $>10^4$ | Yes | *K. pneumoniae* | 4 days |
| 8 | Highly turbid | $>10^4$ | Yes | *E. coli* | 4 days |
| 9 | Turbid | $>10^4$ | Yes | *E. coli* | 3 days |
| 10 | Slightly turbid | $>10^4$ | Yes | *P. aeruginosa* | 3 days |
| 11 | Clear | $<10^4$ | No | - | 2 days |
| 12 | Slightly turbid | $<10^4$ | No | - | 2 days |
| 13 | Slightly turbid | $<10^4$ | No | - | 2 days |

Each sample was processed in duplicate using four different DNA extraction methods: IL-DEx, QIAamp DNA Mini Kit (QIAGEN), MagaZorb DNA Mini-Prep Kit (Promega, USA), and an in-house protocol based on mechanical bead-beating and phenol-chloroform extraction (PC). For each replicate, 1 ml of urine was centrifuged at 10000 x g for 10 minutes at 4 °C, and the pellet was subjected to DNA extraction. Negative extraction controls were included in duplicates for all methods.

Clinical samples were also observed by epifluorescence microscopy. For this, urine samples were diluted 1:10, filtered through 0.2-µm filters (Anodisc 25) and stained with SYBR Gold. Staining and imaging were performed as described above.

### Ionic liquid-based DNA extraction (IL-DEx)

The ionic liquid (IL) [C_2_mim][OAc] (1-ethyl-3-methylimidazolium acetate; purity >95%; CAS 143314-17-4) was purchased from Iolitec (Germany) and used as a 90% (w/w) solution in 10 mM Tris pH 8 buffer. The solution was prepared by weighing the required amount of IL and Tris buffer.

For extraction, 10 µl of the respective sample or the cell pellet were mixed with 90 µl of 90% [C_2_mim][OAc] and incubated at 95 °C for 5 min. The resulting lysate was mixed with 150 µl of SeraSil-Mag™ 400 silica coated superparamagnetic beads (Cytiva, USA). Subsequently, 765 µl of 10 mM Tris pH 8 buffer were added to dilute the lysate and the mixture was vortexed briefly and incubated on a thermomixer (24 °C, up to 10 min, 1400 rpm) to facilitate nucleic acid binding to the beads. Beads were collected by placing the reaction tubes in a magnetic separation rack for 30 - 60 s. After discarding the supernatant, the beads were washed with 500 µl of 70% ethanol prepared in 10 mM Tris pH 8 buffer, then air-dried at room temperature. Elution was performed by adding 100 µl of TE buffer (10 mM Tris-HCl, 1 mM EDTA, pH 8.0) and incubating on a thermomixer (65 °C, 3 min, 1400 rpm).

### Reference extractions

DNA was extracted using the QIAamp DNA Mini Kit (QIAGEN, Germany), following the manufacturer’s protocol for gram-negative or gram-positive bacteria for cultured samples, and the protocol for gram-positive bacteria for spiked urine samples.

For experiments with clinical samples, DNA was extracted with a bead-beating and phenol-chloroform extraction protocol (PC), the QIAamp DNA Mini Kit (QIAGEN), and the MagaZorb DNA Mini-Prep Kit (Promega), both according to the manufacturers’ instructions for gram-positive bacteria. Elution and resuspension volumes for experiments with clinical urine were adjusted to 100 µl for all methods. Extraction with bead-beating and phenol-chloroform was performed as previously described [28-30]. In short, the pellet was resuspended in 0.9% NaCl and cell lysis was achieved by adding CTAB buffer, phenol, chloroform/isoamyl alcohol, and glass beads in a FastPrep 24 bench-top homogeniser (MP Biomedicals Inc., USA) at a speed setting of 6 m/s for 30 s. DNA was precipitated with isopropanol and washed with ethanol. The dried DNA was resuspended in 100 µl of 10 mM Tris pH 8.0 buffer.

### Quantification of DNA using quantitative PCR

Total bacterial DNA was quantified using qPCR targeting the V1-V2 region of the 16S rRNA gene, with primers universally conserved across bacterial taxa (denoted as 16S-qPCR) [31]. *E. coli* DNA was quantified with a qPCR assay targeting the 23S rRNA gene (denoted as *E. coli*-qPCR) [32]. *E. faecalis* and *E. faecium* DNA was quantified with a qPCR assay targeting the *Enterococcus* 23S rRNA gene (denoted as *Enterococcus*-qPCR) [33, 34]. Human DNA was quantified with qPCR targeting the human *Alu* repeats (denoted as human*-*qPCR) [35]. All qPCR reactions were carried out in a total reaction volume of 15 µl containing oligonucleotides, 7.5 µl qPCR Master Mix and 2.5 µl of template DNA. Amplification was performed on a qTOWER^3^ G real-time thermocycler (Analytik Jena, Germany). Detailed information on qPCR assays is provided in Supplementary I Table S1 and S2. Unless otherwise stated, qPCR reactions were carried out in duplicates. Calibration curves were generated using a dilution series of plasmid DNA containing a known number of target gene copies, quantified via PicoGreen measurements. Calibration curves for human-qPCR were generated using a dilution series of human genomic DNA (TaqMan™ Control Genomic DNA, Applied Biosystems, USA) containing a known mass of genomic DNA. To rule out PCR inhibition, samples were measured in multiple dilutions. No template controls (NTCs) were included in each qPCR run.

### Oxford Nanopore full-length 16S rRNA sequencing

The 16S rRNA gene was amplified using modified versions of the 27F and 1492R primers (Supplementary I Table S2), respectively, yielding a near full-length 16S amplicon with an approximate length of 1500 bp. The amplification was performed with the LongAmpTaq HotStart 2x MasterMix (New England Biolabs, USA) in a total reaction volume of 25 µL. The amplicons were purified using the ProNex Size-Selective Purification System (Promega, USA), applying a 1.5-fold ratio of beads-to-sample volume for an approximate size cutoff at 250 bp. The subsequent library preparation was conducted using the Native Barcoding Kit 96 V14 (Oxford Nanopore Technologies, UK), following the manufacturer’s instructions and the recommended amplicon input amount of 200 fmol per sample. The final library was sequenced on a MinION Mk1B platform using an R10.4.1 flow cell (Oxford Nanopore Technologies, UK) for 24 hours. The Nanopore sequencing run and its basecalling were checked with NanoPlot [36]. In case a sample exceeded a yield of 20,000 reads, the raw reads were down-sampled to 20,000 reads using rasusa [37]. Subsequently, the reads were processed with Chopper [36] retaining only those which were between 500 bp and 2,000 bp long. The read statistics were computed with NanoStats [36]. Taxonomic classification was carried out with emu (v3.5.0) [38] using the standard emu database (v.3.4.5; a combination of rrnDB v5.6 [39] and NCBI 16S RefSeq [40]). Read statistics and emu classifications per sample were collected with Python.

### DNA quantity, purity and fragmentation

Total DNA concentration (ng/ml urine) and purity (A260/280 and A260/230 ratios) were measured via absorbance using a NanoDrop One/OneC Spectrophotometer (Thermo Fisher Scientific, USA). DNA fragmentation was assessed by agarose gel electrophoresis. Briefly, a 1% TBE agarose gel was loaded with 10 µl of extracts and 5 or 10 µl of 1 kB Plus DNA Ladder (New England Biolabs, Germany). DNA was stained with SYBR Gold (Thermo Fisher Scientific) and visualized using a Gel Doc XR+ system (Bio-Rad, CA).

### Statistical analysis

Data analysis and visualization were performed using Microsoft Excel and R (version 4.3.3; RStudio). A mixed-design ANOVA was conducted using the ezANOVA() function from the ez package to assess the effects of extraction method (within-subject factor) and sample category (UTI vs. non-UTI; between-subject factor) on bacterial DNA yield. Where the assumption of sphericity was violated (as determined by Mauchly’s test), Greenhouse-Geisser correction was applied. Residuals were tested for normality using the Shapiro-Wilk test (W = 0.99, p = 0.686). Post hoc pairwise comparisons were performed using the emmeans and nlme packages, with p-values adjusted using Tukey’s HSD method. Statistical significance was defined as p < 0.05 (two-tailed). Quantitative PCR (qPCR) data were analyzed using qPCRsoft 4.0 (Analytik Jena). Schematic figures were created with BioRender under academic license j43g432 (Farnleitner, A.).

## Results

### IL-DEx Workflow

The IL-DEx workflow was designed to enable rapid DNA extraction from urine using a single-tube lysis and nucleic acid capture protocol (Fig. 1). The method combines chemical lysis with 1-ethyl-3-methylimidazolium acetate ([C_2_mim][OAc]) and magnetic bead-based recovery. DNA is eluted in less than 30 minutes without the need for hazardous organic solvents. The streamlined setup requires only standard laboratory equipment, including a centrifuge, heating block, and a magnetic rack.

**Fig 1.**
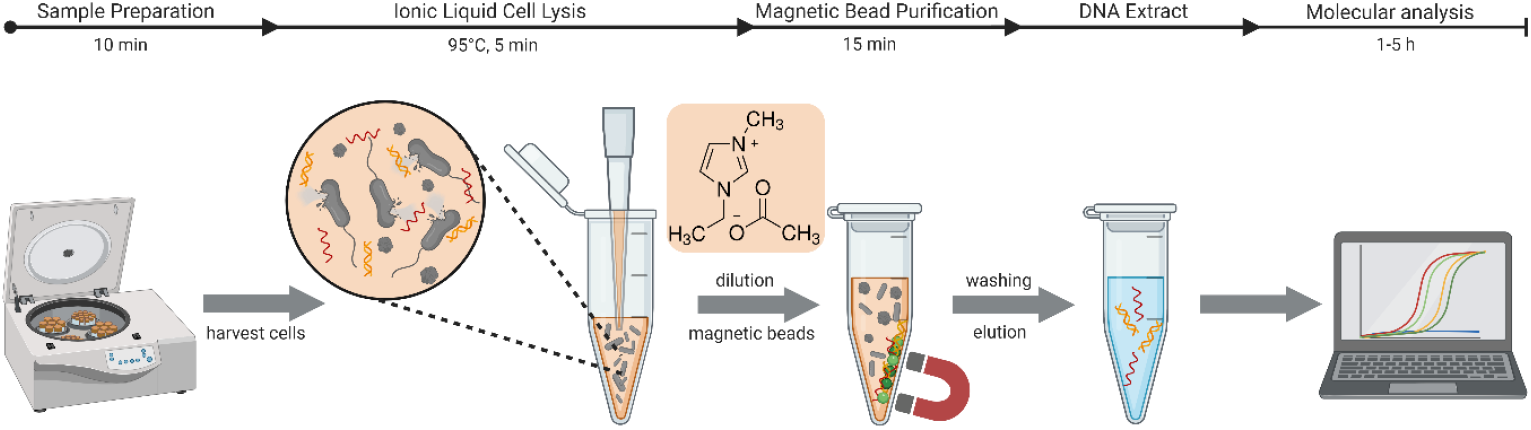
Schematic illustration of the IL-DEx workflow for rapid bacterial DNA extraction from urine samples. The protocol comprises four steps: i) centrifugation to harvest bacterial cells, ii) resuspension of the pellet and ionic liquid-mediated lysis of cells using [C_2_mim][OAc] to release genomic DNA, iii) DNA capture and purification with silica-coated magnetic beads, with [C_2_mim][OAc] also serving as the DNA binding buffer, and iv) elution of purified DNA for v) downstream molecular analyses such as PCR and sequencing. Approximate processing times are indicated above the workflow.

### DNA recovery from bacterial reference strains and clinical isolates

We first evaluated the extraction efficiency of IL-DEx using pure cultures of seven clinically relevant uropathogens: *E. coli, K. pneumoniae, P. aeruginosa, P. mirabilis, E. faecalis, E. faecium*, and *S. saprophyticus*. Fresh cell suspensions were prepared from both a reference strain and a clinical isolate obtained from UTI patients and extracted. DNA yields were determined by quantifying 16S rRNA gene copy numbers, applying a universal qPCR assay (16S-qPCR) across all extracts to minimize assay-specific variability in analytical sensitivity. Extraction efficiencies were calculated by comparing the DNA yields to those obtained using the QIAamp DNA Mini Kit (QIAGEN), which served as a benchmark. The QIAGEN kit was assigned 100% efficiency based on preliminary data showing strong concordance between detected genome copy numbers and bacterial cell counts (Supplementary I, Table S3).

For gram-negative species (*E. coli, K. pneumoniae, P. aeruginosa, P. mirabilis*), IL-DEx yielded DNA amounts comparable to the QIAGEN kit, with relative extraction efficiencies ranging from 47 to 102% (Table 2). No considerable differences were observed between reference strains and clinical isolates. In contrast, extraction efficiencies from gram-positive bacteria were lower, ranging from 0.7% to 8% relative to the QIAGEN kit, likely due to the increased resistance of their thicker cell walls to chemical lysis [41]. Nonetheless, IL-DEx consistently recovered sufficient DNA for downstream analysis, averaging ∼10^7^ 16S rRNA gene copies from 10 µl of cultures with optical densities (OD_600_) near 1. Notably, clinical isolates of *E. faecalis* and *S. saprophyticus* showed higher recoveries (8% and 3%, respectively) than their respective reference strains (0.8% and 0.7%).

**Table 2.** Total bacterial DNA yield (expressed as 16S rRNA gene copies) extracted from gram-negative and gram-positive uropathogens using the IL-DEx protocol (ID-DEx) and the QIAamp DNA Mini Kit (QIAGEN). Full raw data are available in Supplementary II, Table S1.

| Strain<sup>a</sup> | OD<sub>600</sub><sup>b</sup> | Method | Total bacterial DNA extracted<sup>c</sup><br>[Mean 16S rRNA gene copies ± SD] | RSD [%]<sup>d</sup> | Extraction efficiency [%]<sup>e</sup> |
| --- | --- | --- | --- | --- | --- |
| *E. coli*<br>NCTC 9001 | 1.1 | IL-DEx | 1.12 × 10<sup>9</sup> ± 2.12 × 10<sup>8</sup> | 19 | 85 |
|  |  | QIAGEN | 1.32 × 10<sup>9</sup> ± 8.45 × 10<sup>7</sup> | 6 | 100 |
| *E. coli*<br>IHM 5531 | 0.8 | IL-DEx | 3.37 × 10<sup>8</sup> ± 6.93 × 10<sup>7</sup> | 21 | 85 |
|  |  | QIAGEN | 3.95 × 10<sup>8</sup> ± 8.23 × 10<sup>7</sup> | 21 | 100 |
| *P. aeruginosa*<br>NCTC 10662 | 1.0 | IL-DEx | 7.04 × 10<sup>8</sup> ± 1.42E × 10<sup>8</sup> | 20 | 102 |
|  |  | QIAGEN | 6.88 × 10<sup>8</sup> ± 6.56 × 10<sup>7</sup> | 10 | 100 |
| *P. aeruginosa*<br>IHM 5716 | 0.7 | IL-DEx | 3.73 × 10<sup>8</sup> ± 8.69 × 10<sup>7</sup> | 23 | 62 |
|  |  | QIAGEN | 6.03 × 10<sup>8</sup> ± 9.27 × 10<sup>7</sup> | 15 | 100 |
| *K. pneumoniae*<br>DSM 30104 | 1.0 | IL-DEx | 8.49 × 10<sup>8</sup> ± 1.08 × 10<sup>8</sup> | 13 | 56 |
|  |  | QIAGEN | 1.53 × 10<sup>9</sup> ± 1.71 × 10<sup>8</sup> | 11 | 100 |
| *K. pneumoniae*<br>IHM 5571 | 0.8 | IL-DEx | 2.17 × 10<sup>8</sup> ± 5.16 × 10<sup>7</sup> | 24 | 47 |
|  |  | QIAGEN | 4.62 × 10<sup>8</sup> ± 1.67 × 10<sup>7</sup> | 4 | 100 |
| *P. mirabilis*<br>DSM 4479 | 1.1 | IL-DEx | 1.52 × 10<sup>9</sup> ± 2.53 × 10<sup>8</sup> | 17 | 90 |
|  |  | QIAGEN | 1.69 × 10<sup>9</sup> ± 3.02 × 10<sup>8</sup> | 18 | 100 |
| *P. mirabilis*<br>IHM 5533 | 0.7 | IL-DEx | 4.49 × 10<sup>8</sup> ± 5.24 × 10<sup>7</sup> | 12 | 78 |
|  |  | QIAGEN | 5.74 × 10<sup>8</sup> ± 7.87 × 10<sup>7</sup> | 14 | 100 |
| *E. faecalis*<br>DSM 20478 | 1.1 | IL-DEx | 9.31 × 10<sup>6</sup> ± 8.61 × 10<sup>5</sup> | 9 | 0.8 |
|  |  | QIAGEN | 1.23 × 10<sup>9</sup> ± 3.81 × 10<sup>8</sup> | 31 | 100 |
| *E. faecalis*<br>IHM 5490 | 0.7 | IL-DEx | 3.49 × 10<sup>7</sup> ± 1.54 × 10<sup>7</sup> | 44 | 8.3 |
|  |  | QIAGEN | 4.18 × 10<sup>8</sup> ± 8.66 × 10<sup>7</sup> | 21 | 100 |
| *E. faecium*<br>DSM 20477 | 1.3 | IL-DEx | 2.52 × 10<sup>7</sup> ± 2.16 × 10<sup>6</sup> | 9 | 3 |
|  |  | QIAGEN | 8.56 × 10<sup>8</sup> ± 3.07 × 10<sup>8</sup> | 36 | 100 |
| *E. faecium*<br>IHM 5458 | 0.8 | IL-DEx | 8.74 × 10<sup>6</sup> ± 1.68 × 10<sup>6</sup> | 19 | 2 |
|  |  | QIAGEN | 4.22 × 10<sup>8</sup> ± 1.84 × 10<sup>7</sup> | 4 | 100 |
| *S. saprophyticus*<br>DSM 20229 | 1.1 | IL-DEx | 3.89 × 10<sup>6</sup> ± 4.00 × 10<sup>4</sup> | 1 | 0.7 |
|  |  | QIAGEN | 5.31 × 10<sup>8</sup> ± 3.20 × 10<sup>7</sup> | 6 | 100 |
| *S. saprophyticus*<br>IHM 5493 | 0.8 | IL-DEx | 7.53 × 10<sup>6</sup> ± 5.28 × 10<sup>5</sup> | 7 | 3 |
|  |  | QIAGEN | 2.21 × 10<sup>8</sup> ± 4.01 × 10<sup>7</sup> | 18 | 100 |

<sup>a</sup> The panel includes both reference strains and clinical isolates (denoted as “IHM”), all freshly cultured and processed in triplicate. DNA was extracted from 10 µl of bacterial suspension per replicate.
<sup>b</sup> OD<sub>600</sub> refers to the optical density of the bacterial cell suspension used for extraction.
<sup>c</sup> Data are reported as mean 16S rRNA gene copies ± standard deviation (SD) from three biological replicates.
<sup>d</sup> Relative standard deviation (RSD, %) reflects inter-replicate variability.
<sup>e</sup> Extraction efficiency (%) represents the DNA yield recovered with IL-DEx relative to the QIAGEN kit, which was defined as 100%.

### DNA recovery from spiked urine samples

To assess extraction performance in urine matrices, we conducted spiking experiments in i) artificial urine (AU), which lacks background DNA and ii) native urine from healthy donors, which contains both host and microbial DNA.

#### Spiking experiments in artificial urine (AU)

AU experiments allowed direct comparison of extraction efficiency and detection sensitivity between IL-DEx and the QIAamp DNA Mini Kit (QIAGEN) under well-defined conditions. For each of the seven tested uropathogens, serial dilutions of fresh bacterial suspensions were prepared in AU at final concentrations ranging from ∼10^8^ to ∼10^2^ CFU/ml. One-milliliter aliquots were extracted in triplicate using both IL-DEx and the QIAGEN kit. DNA yields were quantified by qPCR targeting the 16S rRNA gene.

As shown in Figure 2, IL-DEx yielded linear DNA recovery across all concentrations and strains tested (R^2^ >0.99). In contrast, the QIAGEN kit showed signs of saturation at higher bacterial loads, particularly for gram-negative species, suggesting column overloading. *P. mirabilis* was the exception, for which both methods remained linear across all concentrations. For gram-negative species, IL-DEx yielded DNA amounts comparable to the QIAamp kit. For gram-positive species, IL-DEx yielded approximately 1-2 log_10_ fewer 16S rRNA gene copies per ml.

**Fig 2.**
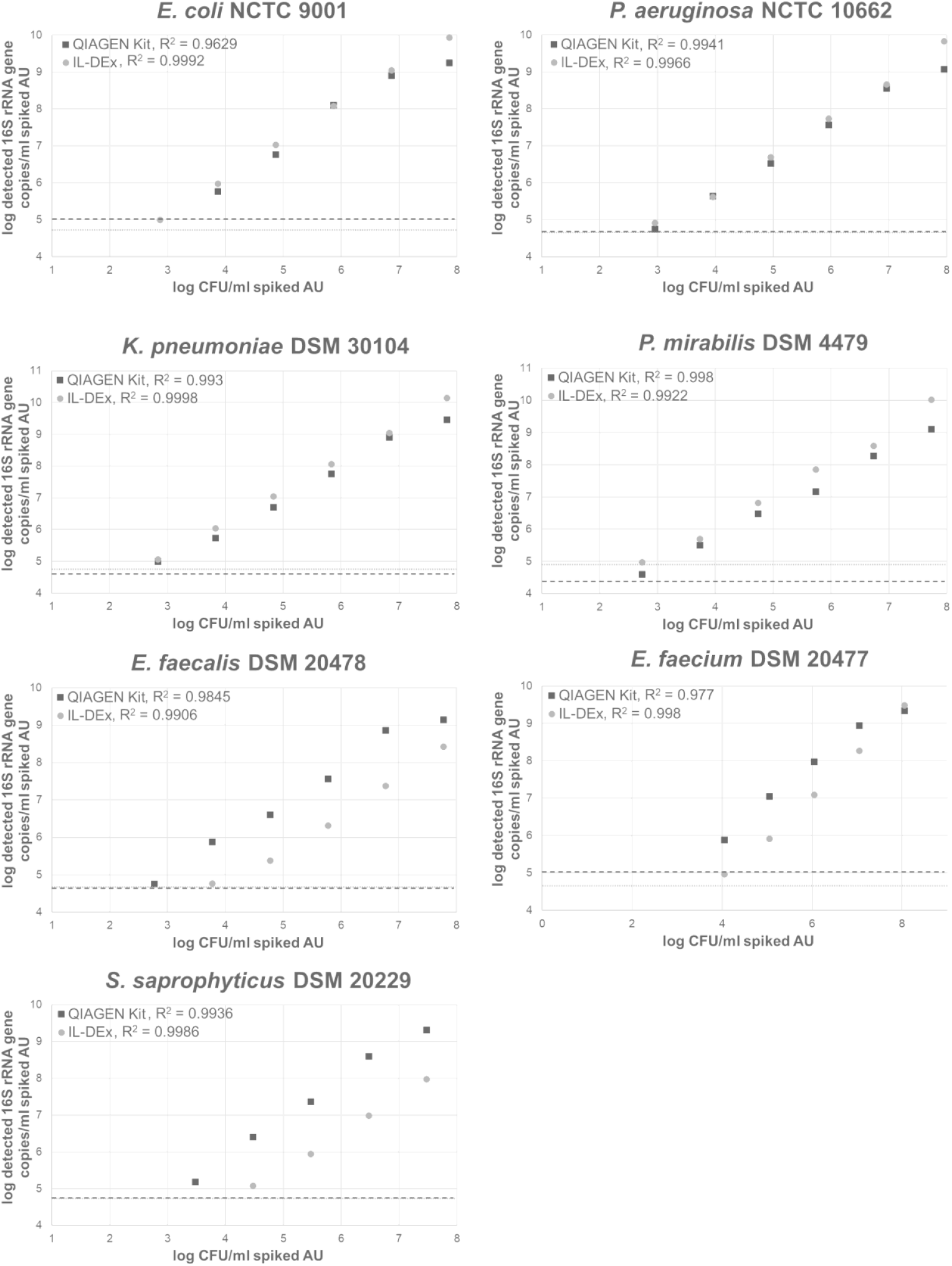
Spiking experiments in artificial urine (AU) to assess linearity and the detection limit of the IL-DEx workflow. Plots show log-transformed 16S rRNA gene copies recovered from 1 ml spiked AU, containing ∼10^8^ to ∼10^2^ CFU/ml. Each dilution was extracted in triplicate with IL-DEx (light grey filled circles) and the QIAGEN kit (dark grey filled squares). Data points represent means. Standard deviations are not shown due to log-scale compression and close overlap between IL-DEx and QIAGEN data, particularly for gram-negative strains; replicate variability is reported in Supplementary II, Tables S2–S8. Dashed and dotted lines indicate background 16S rRNA gene levels detected in extraction controls (unspiked AU) (dashed lines, 16S background extraction control QIAGEN kit; dotted lines, 16S background extraction control IL-DEx). A three-sigma threshold was chosen to exclude all datapoints that lie within three standard deviations of the control mean.

The estimated detection limit for IL-DEx was ∼10^3^ CFU/ml for gram-negatives and ∼10^4^ CFU/ml for gram-positives, based on 16S rRNA gene quantification. However, assay sensitivity was ultimately constrained by low-level background amplification in 16S-qPCR, a phenomenon previously attributed to trace *E. coli* DNA contamination in recombinant polymerases or other laboratory reagents, including extraction kits [42]. This background was evident in extraction controls and defined the detection threshold (dotted and dashed lines in Fig. 2; 3σ cutoff). To improve detection sensitivity, species-specific qPCR assays targeting the 23S rRNA gene were applied for *E. coli, E. faecalis*, and *E. faecium*. These assays confirmed lower detection limits of ∼10^2^ CFU/ml for *E. coli* and ∼10^3^ CFU/ml for the two *Enterococcus* species (Supplementary I, Fig. S1).

#### Spiking experiments in native urine from healthy donors

To simulate sample complexity – including both human host and microbial DNA background – and assess potential matrix effects, urine from three healthy female and two healthy male donors was spiked with uropathogenic *E. coli* (isolate IHM 5531). Preliminary qPCR screening of unspiked urine revealed background 16S rRNA gene levels of ∼10^8^ copies/ml in female donors and ∼10^7^ copies/ml in male donors. Spiking was therefore performed at ∼10^9^ 16S rRNA gene copies/ml (corresponds to 10^7^ *E. coli* CFU/ml urine) to ensure detection above the background. All samples were processed using both IL-DEx and the QIAGEN kit. In addition, unspiked urine was assessed for bacterial and host DNA background by 16S-qPCR and human-qPCR. Both extraction methods enabled robust detection of the *E. coli* spike, with IL-DEx yields comparable to QIAGEN (79% mean extraction efficiency compared to the kit) across donor samples (Fig. 3). IL-DEx underperformed in two unspiked female samples (urine no. 2 and 5), likely reflecting contributions from vaginal flora rich in gram-positive bacteria (e.g., *Lactobacillus*) [43], which are more resistant to lysis. Background DNA analysis further showed that IL-DEx recovered slightly more human DNA than QIAGEN in unspiked urine (334 ng vs. 91 ng; Supplementary I, Fig. S2).

**Fig 3.**
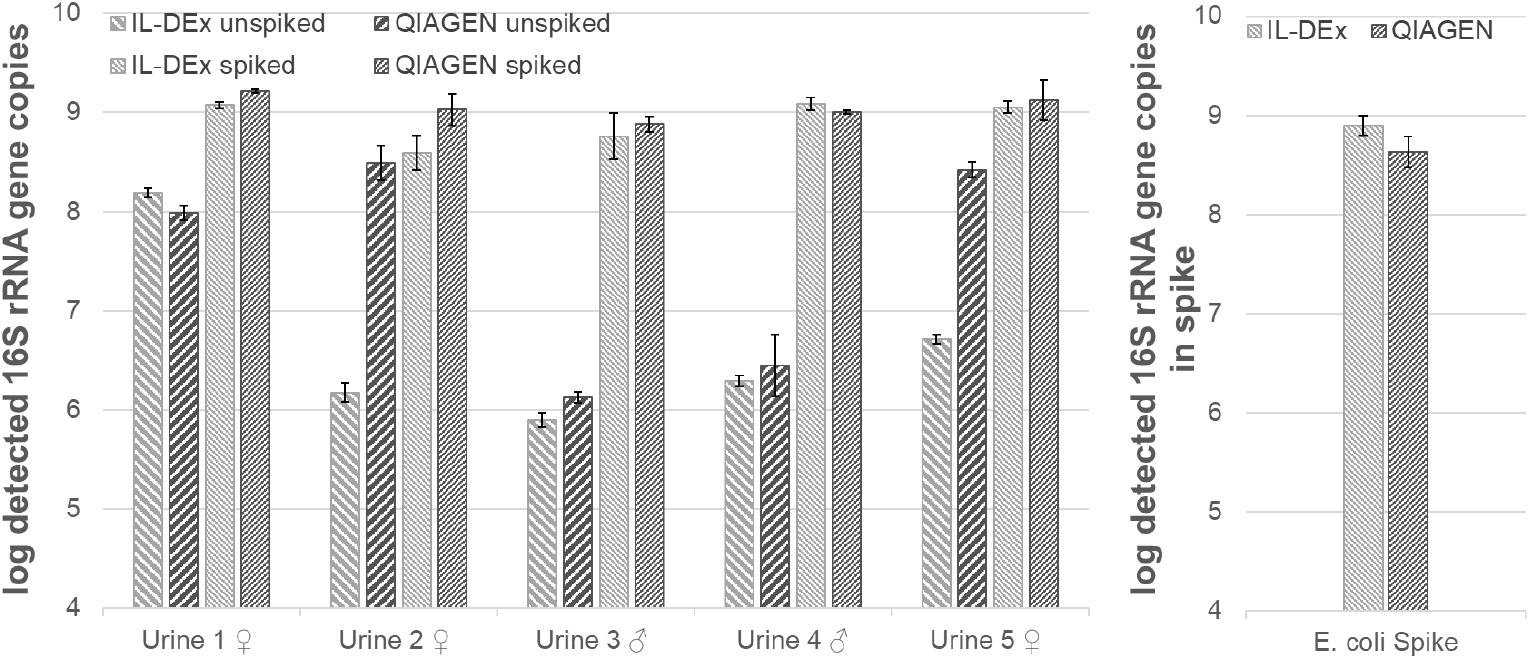
Log-transformed 16S rRNA gene copies detected in extracts from 1 ml spiked and unspiked urine (left panel), as well as from the E. coli spike (right panel). DNA was extracted using IL-DEx (light grey) and the QIAGEN kit (dark grey). Urines were spiked to approx. 10^7^ CFU/ml. Data represent mean values from three biological replicates, whiskers indicate the standard deviation. Raw data are provided in Supplementary II, Table S12.

To further assess gram-positive recovery, a spiking experiment was also performed with *E. faecalis* using urine from a female donor. Samples were processed with IL-DEx and the QIAGEN kit and analyzed by *Enterococcus*-specific 23S qPCR. In unspiked urine, both methods yielded similar background levels. However, in the *E. faecalis* spike and spiked urine sample, IL-DEx recovered ∼1–2 log_10_ fewer *Enterococcus* gene copies than QIAGEN (Supplementary I, Fig. S3). Comparable reductions were also observed in pure culture and artificial urine spiking experiments, indicating that lower yields are primarily due to the intrinsic extraction efficiency for this gram-positive bacterium rather than matrix effects.

### DNA recovery from clinical urine samples

We next evaluated the performance of IL-DEx on 13 clinical urine specimens (10 culture-positive “UTI” and 3 culture-negative “non-UTI” samples as control; Table 1 and Supplementary I, Fig. S4). Each sample was processed using four DNA extraction methods: IL-DEx, two commercial kits (QIAGEN and Promega), and an in-house phenol-chloroform protocol with mechanical bead beating. Extraction performance was broadly assessed by measuring total DNA (via Nanodrop), bacterial DNA (via 16S-qPCR), *E. coli* DNA (via *E. coli*-qPCR), host DNA (via human-qPCR), DNA integrity (via agarose gel) and microbial composition by full length 16S rRNA gene sequencing. All urine samples were also imaged by epifluorescence microscopy (EFM).

#### Total DNA yield and purity

Among all methods, the Promega kit produced the highest total DNA amounts, followed by the phenol-chloroform extraction, IL-DEx and QIAGEN (Fig. 4A). Purity ratios (A260/280) were comparable across methods and found within the acceptable range of 1.8-2.0, whereas A260/230 ratios were lower for IL-DEx (Fig. 4B). Gel electrophoresis of DNA extracts revealed intact high molecular weight DNA in IL-DEx and QIAGEN extracts, while DNA from Promega kit and phenol-chloroform extractions appeared more fragmented (Supplementary I, Fig. S5).

**Fig 4.**
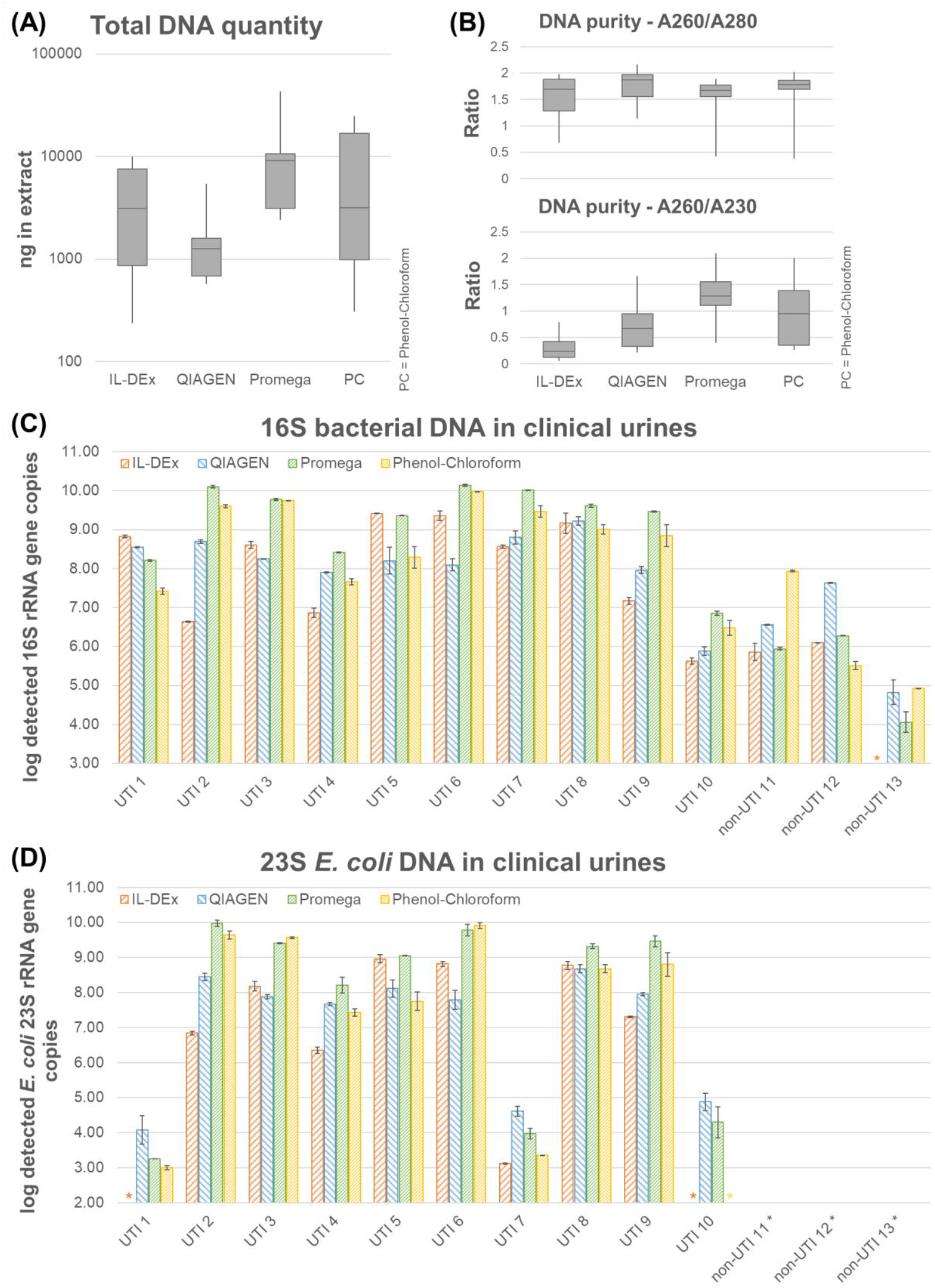
(A) Boxplots of total DNA content (as ng) of all extracts from each method, measured via absorbance. (B) Boxplots of DNA purity (absorbance rates) of all extracts from each method. (C) and (D) Content of total bacterial DNA (as log detected 16S rRNA gene copies) and E. coli DNA (as log detected E coli 23S rRNA gene copies) in the DNA extracts from clinical urines obtained via IL-DEx (orange), QIAGEN kit (blue), Promega kit (green) and phenol-chloroform extraction (yellow). Data shown are mean values from two biological replicates, whiskers indicate the two measured values. For each replicate 1 ml of urine was processed. Asterisks indicate results that lie within three standard deviations from the mean of the extraction controls and were therefore excluded. Full raw data are available in Supplementary II, Table S15-S17.

#### Bacterial DNA Recovery

Bacterial DNA yields (16S rRNA gene copies, Fig. 4C) corresponded well with bacterial cell loads observed by epifluorescence microscopy (Supplementary I, Fig. S6). Non-UTI samples and UTI sample no. 10 (low bacterial cell loads) yielded the lowest 16S rRNA gene copies. A mixed-design ANOVA confirmed significant effects of sample category (UTI vs. non-UTI, F_1,24_ = 36.35, p < 0.0001), extraction method (F_3,72_ = 3.43, p = 0.021), and their interaction (F_3,72_ = 6.62, p = 0.0005). Among UTI samples, Promega and phenol-chloroform yielded significantly more bacterial DNA than IL-DEx (*p* < 0.0001 and *p* = 0.0228, respectively), while IL-DEx and QIAGEN did not differ (*p* = 0.93). For non-UTI samples, no significant differences between methods were detected (all *p* > 0.1). Pairwise comparisons further showed IL-DEx yielded significantly more bacterial DNA in UTI than in non-UTI samples (Δ = 5.9 log_10_ gene copies, p = 0.0001).

#### E. coli-specific DNA detection

*E. coli*-qPCR confirmed the presence of *E. coli* in all culture-positive *E. coli* urine samples (UTI sample no. 2-6, 8, and 9; Fig. 4D and Table 1). In these samples, 23S rRNA gene copy numbers closely matched total bacterial DNA levels (16S rRNA gene copies, Fig. 4C). Low-level signals were also detected in samples 1, 7, and 10.

#### Host DNA Recovery

Human DNA recovery mirrored bacterial DNA extraction trends, where extraction methods with higher bacterial DNA yields (Promega, phenol-chloroform) also extracting higher amounts of host DNA (Supplementary Fig. S7).

#### Microbial profiling via Nanopore Sequencing

Full-length 16S sequencing confirmed that all four extraction methods enabled detection of the dominant uropathogen identified by clinical culture (Fig. 5). In IL-DEx extracts, the dominant pathogen was detected at ≥75% relative abundance, except in UTI sample 10. In this sample, IL-DEx extract sequencing identified *E. coli*, whereas urine culture and sequencing of extracts from QIAGEN, Promega and phenol-chloroform methods identified *P. aeruginosa*. However, *E. coli*-specific qPCR showed no detection in the IL-DEx extract (Figure 4C), while a subsequent qPCR assay specific for *P. aeruginosa* confirmed its presence (data not shown), suggesting a possible sample mix-up during NGS library preparation. Aside from this discrepancy, the microbial community profiles were consistent across methods in UTI samples. Despite known challenges in lysing Gram-positive bacteria, IL-DEx still recovered detectable levels of these taxa (e.g. *Enterococcus, Streptococcus, Lactobacillus, Staphylococcus, Gardnerella* sp.; Fig. 5).

**Fig 5.**
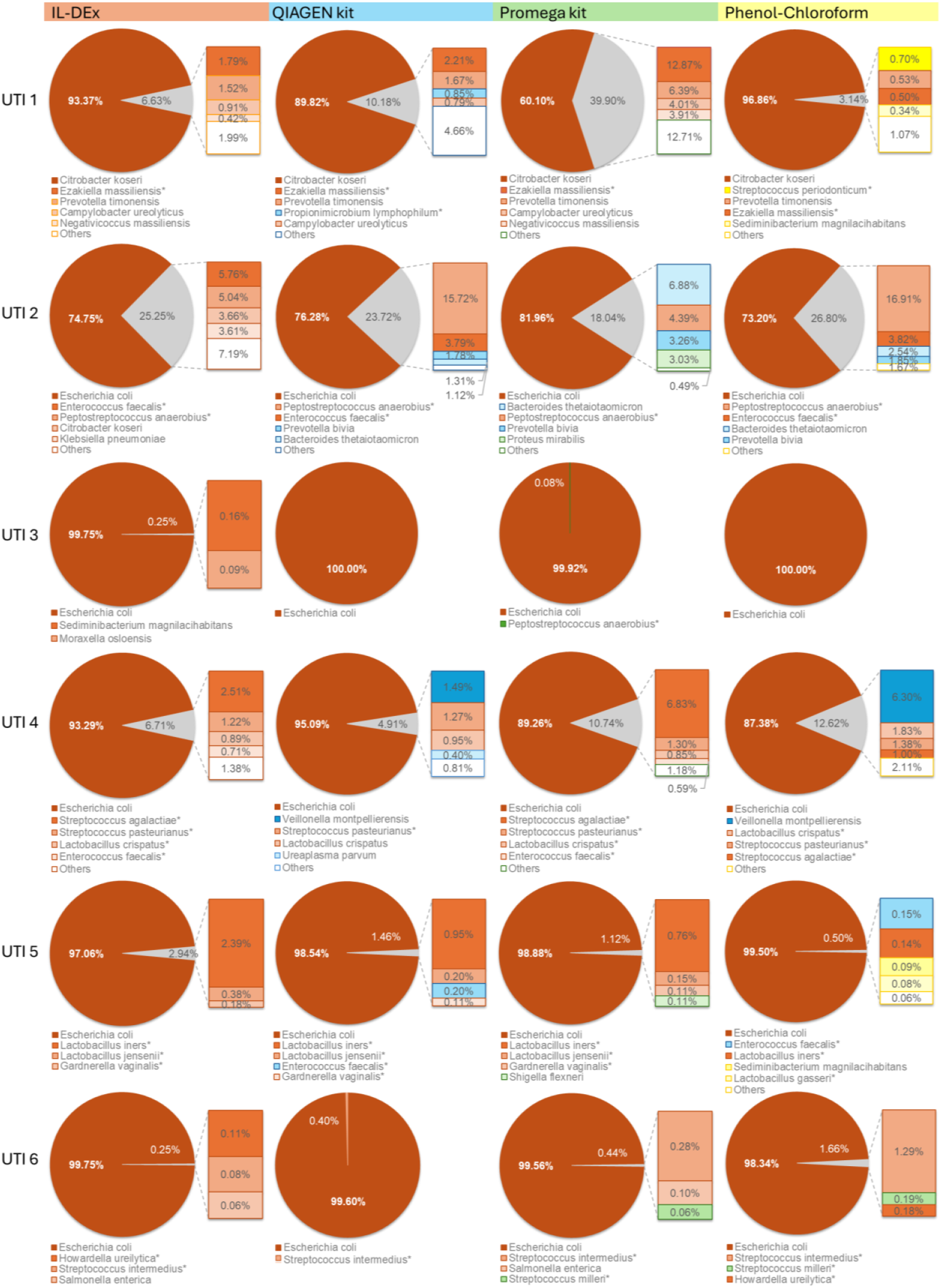
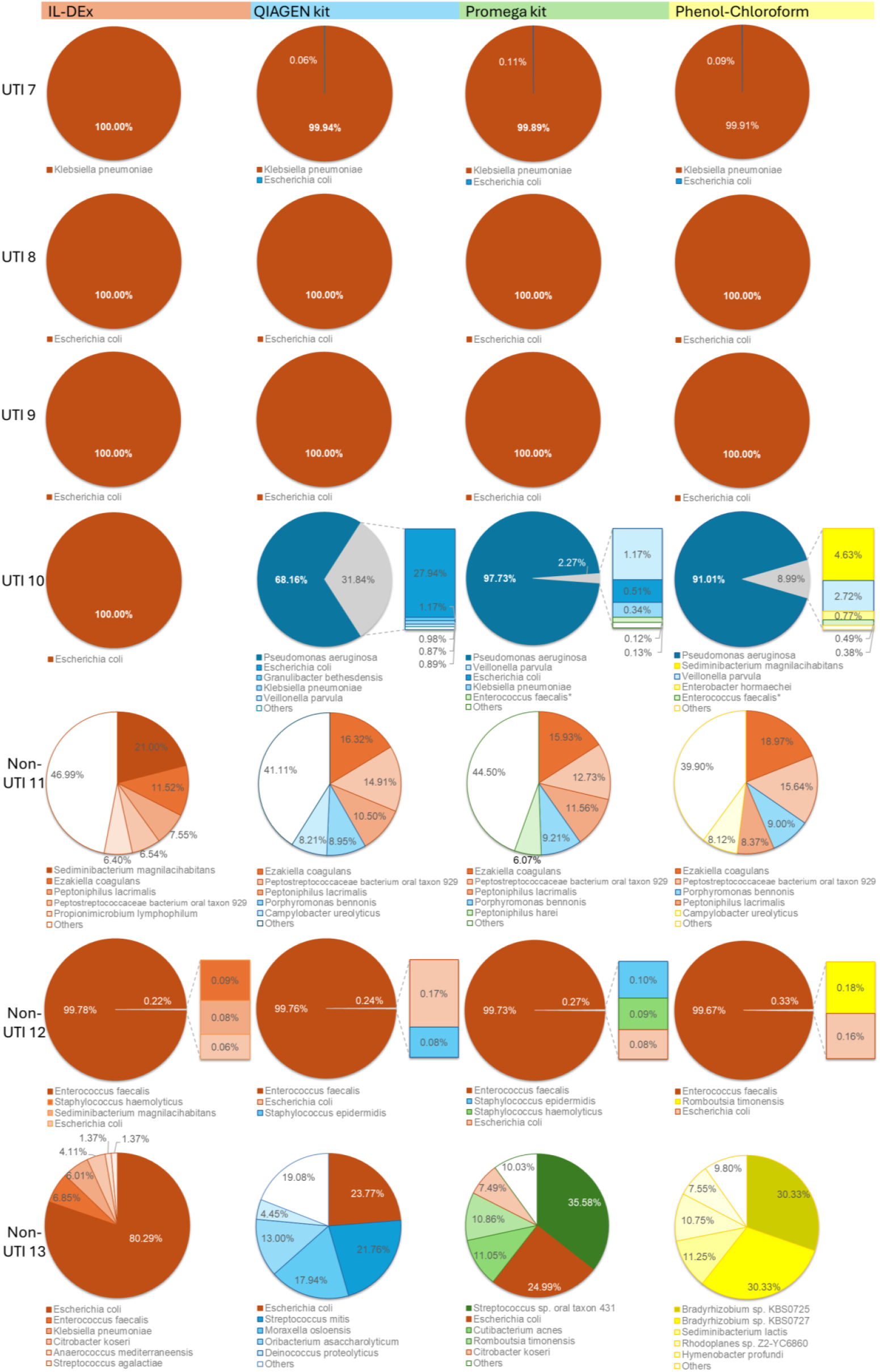
Full length 16S rRNA gene sequencing results for the extracts from the ten UTI and three non-UTI samples obtained via IL-DEx, QIAGEN kit, Promega kit and phenol-chloroform extraction. The five most abundant species are shown. Asterisks indicate gram-positive bacteria. Full raw data are available in Supplementary II Table S19.

## Discussion

Rapid and reliable identification of uropathogens is essential for effective management of urinary tract infections (UTIs). With rising diagnostic demand and increasing antimicrobial resistance, molecular techniques such as PCR, isothermal amplification, and deep sequencing are gaining traction to expand current diagnostic options for UTIs [18, 44]. However, their sensitivity, specificity, turnaround time and cost are impacted by upstream DNA extraction steps. Extraction biases affecting microbial composition are well recognized, underscoring the importance of selecting suitable extraction methods for different sample types [45-47].

In this study, we demonstrated the feasibility of using ionic liquid chemistry for direct DNA extraction from urine, a specimen type that remains technically challenging for molecular diagnostics. Urine is characterized by variable microbial loads, abundant human host DNA, and inhibitory compounds (e.g., urea, salts, calcium oxalate crystals) that can impede bacterial DNA recovery [48-50]. Although recent studies have optimized or compared different extraction approaches for urinary microbiota analyses, alternatives beyond traditional enzymatic, chemical or mechanical disruption remain largely unexplored. Against this background, we evaluated IL-DEx, an ionic liquid-based method that integrates bacterial lysis and nucleic acid capture in a single-tube workflow using the hydrophilic ionic liquid [C_2_mim][OAc] and silica-coated magnetic beads. Compared to commercial kits and a phenol-chloroform protocol tested, IL-DEx offered several advantages: faster processing (∼20 min), lower reagent costs (∼2 € per prep), and elimination of hazardous solvents and specialized homogenization equipment (Supplementary I, Table S4). While not entirely instrument free – requiring centrifugation and heating for cell harvest and lysis – its simplicity and reagent accessibility support its application in both centralized laboratories and decentralized diagnostic settings.

Across spiking experiments and clinical specimens, IL-DEx produced DNA yields for gram-negative uropathogens comparable to the QIAamp DNA Mini Kit (equivalent to the DNeasy Blood and Tissue Kit, QIAGEN), which prior benchmarking studies identified as one the most reliable options for urinary microbiota extraction [26, 51, 52]. Matching QIAGEN’s performance for gram-negatives is clinically significant given that *Escherichia coli, Klebsiella pneumoniae, Proteus mirabilis*, and *Pseudomonas aeruginosa* cause the majority of community- and hospital-acquired UTIs [9, 10]. In contrast, IL-DEx consistently yielded lower recovery from gram-positive bacteria (∼ 1 log_10_ lower), consistent with studies reporting that thick-walled organisms require more rigorous lysis [41, 53]. Recovery for gram-positive bacteria could be increased by centrifuging a larger urine volume for the extraction. Nevertheless, the recovered DNA was sufficient for downstream qPCR and long-read 16S sequencing. Sequencing of DNA extracts confirmed the presence of several gram-positive taxa at low abundance, in agreement with other tested extraction methods and alongside dominant uropathogens (Fig. 5). These included *Lactobacillus spp*. (UTI 5), *Streptococcus spp*. (UTI 4), *Enterococcus spp*. (UTI 2), *Gardnerella vaginalis* (UTI 5), *Ezakiella massiliensis* (UTI 1), and *Peptostreptococcus anaerobius* (UTI 2). Together, these findings demonstrate IL-DEx’s capacity to capture diverse microbial profiles, though clinical interpretation of non-dominant taxa was beyond the scope of this study.

IL-DEx is therefore competitive with commercial silica spin-column methods for gram-negatives (QIAGEN), while offering a faster and chemically safe workflow. In comparison, Promega and phenol-chloroform protocols produced higher total DNA yields, but with more fragmented DNA and substantially more complex procedures, respectively. This trade-off echoes observations of earlier comparative studies, where yield advantages did not always translate into improved sequencing outcomes [26, 54]. Thus, the combination of rapid processing, recovery of high molecular weight DNA, and simplified chemistry distinguishes IL-DEx from conventional extraction methods.

While the results of this study are encouraging, several limitations must be acknowledged. First, clinical evaluation was performed on a small cohort of 13 urine specimens (10 culture-positive, 3 culture-negative), limiting conclusions about diagnostic sensitivity and specificity across diverse patient populations. Larger studies across patient groups and clinical presentations, with clearly defined downstream applications (e.g. multiplex PCR) are needed to establish clinical robustness. Second, although IL-DEx demonstrated compatibility with qPCR and nanopore-based 16S sequencing, its integration with other molecular platforms, such as isothermal amplification or shotgun metagenomic workflows remains to be evaluated. Human-to-bacteria DNA ratios varied between samples (1:1 to 1000:1), as determined by qPCR of 16S and human DNA content. Such variability has minimal impact on amplification-based assays (PCR, isothermal methods, targeted sequencing) but may impact microbial sequencing depth and sensitivity in untargeted (shotgun metagenomics) sequencing approaches [4]. Third, fungal pathogens were not assessed in this study. Although less frequent than bacterial UTIs, *Candida* spp. are commonly implicated in catheter-associated UTIs and are associated with adverse outcomes in immunocompromised or critically ill patients [55]. Given their chitin-rich cell walls, effective DNA extraction may require additional lysis strategies. Future work should explore IL-DEx’s utility in fungal diagnostics and polymicrobial infections.

In conclusion, IL-DEx offers a rapid, simple, and cost-effective approach to DNA extraction from urine specimens. Its compatibility with molecular platforms, ability to recover DNA from clinically relevant uropathogens, and potential for automation support its use in next-generation diagnostic workflows. As molecular testing continues to shift toward rapid, near-patient, and point-of-care applications, streamlined methods such as IL-DEx can help overcome pre-analytical barriers and extend access to advanced infectious disease diagnostics.

## Acknowledgements

This work was funded by the Gesellschaft für Forschungsförderung NÖ (GFF) under the RaCeDeLys project (FTI21-D-012).

## Author contribution

J.K. and C.K. designed the study and wrote the manuscript. J.K. conducted the experiments and analyzed the data, with support of C.K., K.P., R.M., and M.A. B.S., I.J.P., and D.W. provided clinical expertise as well as isolates and specimens. C.K. supervised the project. G.R.H helped supervise the project. G.R.H. and A.H. secured funding. G.R.H., B.S., I.J.P., A.H.F. provided critical feedback and helped shape the manuscript. All authors reviewed and approved the final version of the manuscript.

## Data availability

Oxford nanopore sequencing data have been deposited in the European Nucleotide Archive (ENA) under accession numbers ERS26910746–ERS26910803, associated with project PRJEB97800. Raw data from qPCR analyses can be found in the supplementary files (Supplementary Material II).

